# Long-lasting event-related beta synchronizations of electroencephalographic activity in response to support-surface perturbations during upright stance

**DOI:** 10.1101/2020.06.07.138461

**Authors:** Akihiro Nakamura, Yasuyuki Suzuki, Matija Milosevic, Taishin Nomura

**Author notes:** Corresponding Author: Taishin Nomura, Ph.D., Division of Mechanical Science and Bioengineering, Graduate School of Engineering Science, Osaka University, Osaka 560-8531, Japan.

## Abstract

Movement related beta band cortical oscillations, including beta rebound after execution and/or suppression of movement, have drawn attention in upper extremity motor control literature. However, fewer study focused on beta band oscillations during postural control in upright stance. Here, we examined beta rebound and other components of electroencephalogram (EEG) activity during perturbed upright stance to investigate supraspinal contributions to postural stabilization. Particularly, we aimed to clarify the timing and duration of beta rebound within a non-sustained, but long-lasting, postural recovery process that occurs more slowly compared to upper extremities. To this end, EEG signals were acquired from nine healthy young adults in response to a support-surface perturbation, together with the center of pressure (CoP) and mass (CoM) and electromyogram (EMG) activities of ankle muscles. Event-related potentials (ERPs) and event-related spectral perturbations were computed from EEG data using the perturbation-onset as a triggering event. After short-latency (< 0.3 s) ERPs, our results showed high-beta band power decrease (event-related desynchronization), which was followed by an event-related synchronization at high-beta band and theta band desynchronization. Specifically, beta synchronization (beta rebound) was sustained for as long as three seconds. EMGs of the ankle muscles and the ankle and hip joint torques remained activated in the first half period of the beta rebound. They returned to the steady-state in the remaining phase, where the CoP/CoM were in their final approach to the equilibrium. We propose possible mechanistic causes of the long-lasting beta rebound, which may be related to underlying intermittent control strategy in upright stance.

**New & Noteworthy:** Beta rebound cortical activity was identified during postural recovery from a perturbed upright stance. Contrary to upper extremities, it was initiated before the recovery of motion was completed, and sustained for as long as three seconds. Those novel characteristics of the beta rebound might be caused by slow dynamics of the upright posture and by selections of on/off switching in an intermittent feedback controller, which was shown to stabilize upright posture.

## Introduction

Supraspinal contributions to postural stabilization during human upright stance have been demonstrated by postural instability in patients with neurological disorders such as Parkinson’s disease (PD), multiple sclerosis, and stroke (Horak et al. 1992; Yamamoto et al. 2011; Perera et al. 2018; Visser et al. 2008; Frzovic et al. 2000; Geurts et al. 2005). Alterations in patterns of postural sway and postural responses to perturbations induced by a cognitive load (Woollacott and Shumway-Cook 2002) and by motor learning during adapting to novel environmental demands (Horak and Nashner 1986) also provide a glimpse into the roles played by the supraspinal networks in the control of upright posture.

Electroencephalogram (EEG) activity and/or motoneuronal responses to transcranial magnetic stimulation (TMS) during standing are more direct methods to characterize electrical activity of the cerebral cortex associated with postural control. For example, using EEG recordings, it has been shown that postural reactions against mechanical perturbations during voluntary postural sway are triggered by the central command mechanisms associated with bursts of gamma-band activities (Slobounov et al. 2005). Moreover, soleus muscle showed larger motor evoked potentials in response to TMS stimuli during standing on a moving planform, compared to standing on a still platform, which implies increased corticospinal excitability in more challenging postural environments (Solopova et al. 2003). Yet, there is relatively limited knowledge on how activities of the cerebral cortex, particularly those measured by EEG, encode sensory information processing and motor control during stabilization of upright stance (Bolton 2015; Jacobs and Horak 2007; Wittenberg et al. 2017).

EEG signals during upright stance have traditionally been investigated using event-related potentials (ERPs) in response to brief postural perturbations, where a perturbation-event-locked average of EEG time series is computed to achieve high signal-to-noise ratio. Particularly, the most representative ERP identified in EEG activities in response to brief mechanical perturbations has a negative potential and it is referred to as the N1 (Quant et al. 2005; Varghese et al. 2017). The N1 potential is typically induced with a latency of about 90-170 ms and amplitudes ranging from −10 to −70 μV, while the responses are spatially distributed over the frontal, central, and parietal cortices (Varghese et al. 2017). There are several interpretations for the origins of the N1 responses. One considers that N1 represents neural processing of sensory information (Dietz et al. 1984; Dimitrov et al. 1996; Staines et al. 2001) necessary for coordinating reactive balance responses based on the fact that the latency and amplitude of N1 are altered by the afferent transmission delay (Dietz et al. 1985) and the level of cognitive load (Quant et al. 2004). Another interpretation is that N1 represents an error signal for detecting postural instability (Adkin et al. 2006, 2008; Mochizuki et al. 2009; Payne et al. 2019). Payne *et al*. have been working to characterize the N1 and other ERPs during perturbed stance in recent years. They showed that: (1) people with lower balance ability exhibited larger N1 responses compared to those with better balance control (Payne and Ting 2020a); and (2) N1 includes startle responses in addition to balance-correcting motor responses (Payne et al. 2018). In addition, they also showed abnormal N2 activity in PD patients (Payne and Ting 2020b), where N2 is induced with a latency of about 400 ms with much smaller amplitudes compared to N1, suggesting that later components might be less reflexive and more supraspinal-network-dependent. Taken together these studies provide insights into specific cortical responses to balance perturbations during upright stance.

Time-frequency characteristics of EEG responses induced by mechanical perturbations can also be analyzed by computing event-related spectral perturbations (ERSPs) (Mierau et al. 2017; Peterson and Ferris 2018; Slobounov et al. 2005; Solis-Escalante et al. 2019; Varghese et al. 2014, 2015). Specifically, ERSP represents neural oscillations at distinct frequency-bands to characterize the balance between neuronal excitation and inhibition (Buzsáki et al. 2012; Muthukumaraswamy 2014). Using EEG ERSP, it may be possible to identify cortical responses to particular events during standing as shown during other motor and cognitive tasks (Makeig 1993; Pfurtscheller and Lopes da Silva 1999). For motor tasks other than the upright posture, it is considered that low-frequency band-limited cortical synchronizations (< 20 Hz) are associated with a deactivated state of the corresponding networks, and that high-frequency band-limited cortical synchronizations (> 20 Hz) reflect a state of active information processing (Pfurtscheller and Lopes da Silva 1999). Particularly, event-related desynchronization (ERD) at beta band (13-30 Hz) prior to and/or during motor execution as well as event-related synchronization (ERS) of beta band oscillations after the movement are well known (Pfurtscheller et al. 1996; Pfurtscheller and Lopes da Silva 1999). Beta band ERS is typically referred to as the post-movement rebound or simply as beta rebound, and it is considered to be associated with *status quo* in terms of postural maintenance of upper extremities (Engel and Fries 2010) and afferent sensory information processing (Cassim et al. 2001). Attenuation of the beta rebound in PD patients is consistent with impaired sensory integration (Vinding et al. 2019). Moreover, beta rebound is typically observed in Go/NoGo tasks, with and without motor executions, which is also a typical movement-related EEG response characteristic (Alegre et al. 2004; Swann et al. 2009; Zhang et al. 2008). Since beta ERS for the NoGo response is not accompanied with an actual motor execution, it may represent motor-related decision-making processing in the supraspinal networks, in addition to information processing of sensory feedback signals. Therefore, considering attenuated beta ERS responses reported for the Go/NoGo task in PD patients (Wu et al. 2019), the beta ERD and the subsequent beta ERS (beta rebound) represent information processing performed by the cortico-basal ganglia motor loop.

Among a small number of studies analyzing ERSPs during upright stance, Varghese *et al*. reported that whole-body perturbations during upright stance caused ERS in theta (3-7 Hz), alpha (8-12 Hz), and beta bands, within very short response latencies (at most a few hundred milliseconds), which correspond to the time range of the N1 potentials (Varghese et al. 2014). In addition, some studies showed that brief mechanical perturbations applied by pulling the body during standing posture induced beta ERD after the theta and alpha ERSs with short latencies (Peterson and Ferris 2018). However, beta rebound, which has been examined extensively during voluntary and passive motor tasks in upper extremities, has not been investigated during postural control tasks in upright stance. One of the key issues in dealing with postural recovery processes during upright standing in response to external perturbations is the difference in the time-scale of mechanical dynamics compared to movements of the upper extremities. Namely, postural responses of musculo-skeletal system during upright stance typically persist over longer periods of time (i.e., a few seconds), unlike those of the upper extremities that are typically completed within a few hundred milliseconds.

In this study, we therefore examined ERSPs during the long-lasting postural biomechanical recovery process of upright equilibrium in response to small impulsive (step-like) support-surface perturbations. A particular interest was to investigate the timing and the duration of beta band ERD and ERS responses, and to quantify them. Because a postural recovery process from a perturbed posture to the upright equilibrium is transient, and it settles down eventually to the post-recovery equilibrium state, this could be regarded as a *status quo* for postural maintenance. Therefore, similar to the upper limbs (Engel and Fries 2010), we hypothesized that beta ERD and subsequent post-movement beta rebound (beta ERS) would be present in the EEG cortical activities following upright stance perturbations. If we could observe beta ERD and a beta rebound as hypothesized, the next objective would be to clarify in the temporal profiles of the appearance of beta activities within the long-lasting biomechanical postural response. In the simplest possible situation, beta ERD would appear persistently during the postural response while the muscles are active, and then a beta rebound would appear following postural recovery, i.e., as a post-movement rebound after a few seconds required for the upright posture to be fully recovered. This scenario is based on beta ERD and ERS during upper extremity tasks (Engel and Fries 2010), with possible variations in time intervals due to the longer-lasting postural response. However, if we observe beta ERD and ERS with qualitative differences in temporal characterizations from those during upper extremity tasks, they could lead to shedding new light on mechanistic causes of the beta rebound.

## Methods

### Participants

Nine healthy young male participants (mean age 24.2 years, SD 1.4 years) were included in the study. None of the participants suffered from neurological disorders nor used medications that could influence posture. All participants gave written informed consent, which was executed in accordance with the principles of the Declaration of Helsinki (2013) and approved by the ethical committee of the Graduate School of Engineering Science at Osaka University.

### Experimental protocol

Standing posture, electromyography (EMG), and EEG signals were measured under the following two conditions: quiet standing (control) and support-surface perturbation (perturbed). Postural sway during quiet stance in the control condition was measured to determine the equilibrium posture to define a transient postural recovery from the equilibrium state. In each control or perturbed condition trial, participants were asked to stand still for 7 min on a treadmill (Bertec, Ohio, USA) with their arms folded on their chest, while keeping their gaze fixed on a target located 4.5 m in front of them at the eye level. Specifically, the anterior-posterior (AP) direction of postural sway was defined to be parallel to the longitudinal direction of the treadmill belt. Although 7 min trials are relatively long, it has previously been shown that important sway metrics, including diffusion coefficients of the stabilogram at long and short term regions, are not affected by long standing durations, suggesting that cortical activities associated with postural control are also not affected by long standing durations (e.g., van der Kooij et al. 2011). The participants performed two trials for each of the control and perturbed conditions (four 7 min trials in total). The order of four trials was chosen randomly from C-P-C-P, C-P-P-C, P-C-P-C and P-C-C-P with equal probability, where C and P represent the control and the perturbed trials, respectively. A rest of at least 3 min was given between trials. During the control trials, the participants were asked to remain relaxed and still for the duration of each trail. During the perturbed trials, the support-surface perturbation was applied during upright stance by moving the treadmill-belt backwards slightly and quickly using an in-house computer-program. Specifically, each perturbation spanned 200 ms, which was composed of an acceleration phase (4 m/s^2^ for 100 ms), followed by a deceleration phase (−4 m/s^2^ for 100 ms). In each 7 min trial, 20 perturbations were applied with a fixed interval of 20 s between each perturbation. All participants could maintain upright stance against each perturbation without having to initiate compensatory steps or other overt movements in any of the trails. We confirmed that no participants could make an accurate prediction to brace for the perturbation, based on the following question administered orally at the end of the experiment: “Were you able to predict the occurrence of each perturbation? Please provide yes-no answers.” In all trails, participants were instructed to stand upright in a relaxed state to reduce the effects of muscle-activity-derived artifacts in EEG recordings.

### Experimental setup

Postural kinematics were measured using a three-dimensional optical motion capture system (SMART-DX, BTS Bioengineering, Milan, Italy) with a sampling frequency of 300 Hz, where light reflection markers were attached on ankles, greater trochanters, and acromions of the left and right sides of the body. The postural kinematics in the AP and medio-lateral (ML) directions were quantified by measuring the center of pressure (CoP), for which a force plate built in the treadmill acquired time-profiles of the ground reaction force vectors at a sampling frequency of 1,200 Hz. EMG signals were recorded from the ankle muscles of both the left and the right legs, including the soleus (SO), medial-gastrocnemius (MG) and tibialis anterior (TA) muscles. EMG data were recorded using wireless surface electromyograms (FreeEMG, BTS Bioengineering, Milan, Italy) with a sampling frequency of 1,000 Hz. EEG signals were measured using a 32 channel mobile bio-amplifier that was placed inside a backpack and worn by the participants and a waveguard cap that included active shielded cables for reducing movement-induced interference (eegosports, ANT Neuro, Hengelo, Netherlands). Specifically, Ag/AgCl electrodes were arranged in accordance to the International 10/10 system (Chatrian et al. 1985; Jurcak et al. 2007). All electrodes used CPz as a reference, and one frontal electrode was used as the ground (GND). The impedance in all electrodes was controlled to be less than 20 kΩ during the measurements (Ferree et al. 2001). All EEG signals were sampled at a sampling frequency of 2,048 Hz and stored on a computer for post-processing.

### CoP analysis during quiet stance

Postural sway data during quiet stance (control condition) were characterized using the CoP time-series for each participant. In short, the following parameters were computed: (i) standard deviations of CoP fluctuations in the AP and ML directions; and (ii) slopes of linear regression lines for the log-log plotted power spectrum of CoP in the AP direction at low (0.02-0.2 Hz) and high (1-8 Hz) frequency regimes, as described elsewhere (Matsuda et al. 2016; Yamamoto et al. 2011, 2015). In particular, it is known that the power spectrum in the low frequency regime exhibits the *f* ^-*β*^-type scaling with the exponent close to *β* = 1.5, which is one of the hallmarks of the intermittent postural control hypothesis (Asai et al. 2009; Collins and De Luca 1994; Nomura et al. 2013; Yamamoto et al. 2015). Parameters defined here were computed from the entire 7 min CoP time series and the results obtained on the two control trials were averaged to obtain the CoP postural sway measures for each participant.

### Estimation of Joint angles and CoM for perturbed stance

Joint angles and position of the total body center of mass (CoM) were estimated only for the perturbed stance. To this end, human body during upright stance was modeled using a triple-inverted pendulum consisting of rigid links, with the head-arm-trunk (HAT) link, a leg-link, and a foot-link, moving in the sagittal plane around two pinned-joints corresponding to the ankle and hip joints (Suzuki et al. 2012). See Fig. S1 in Supplemental materials. The foot-link was assumed to be fixed on the support-surface. The ankle joint angle *θ*_a_ was defined as the tilt angle of the leg-link from the vertical upward line. The hip joint angle *θ*_h_ was defined as the relative joint angle between the HAT- and the leg-links. Plantar flexion for the ankle joint and extension for the hip joint were defined as the positive direction. *θ*_a_ and *θ*_h_ for each participant were estimated using the motion-captured positions of the markers in the global coordinate system. Specifically, the positions of two corresponding markers on the left and right sides of the body were projected on the sagittal plane, and then averaged to obtain the position of each of the ankle, greater trochanter, and acromion in the model. For full details related to the model and joint angles estimation, see sections S1 and S2 in Supplementary materials.

Horizontal position of the total body center of mass (CoM) in the AP direction during standing was estimated from the CoM-positions and the masses of the HAT-link (*m*_HAT_) and the leg-link (*m*_L_) of the model. *m*_HAT_ and *m*_L_ were estimated from the total body weight of each participant using a statistical formula of *m*_HAT_: *m*_L_ = 0.62: 0.35 (Suzuki et al. 2012). We assumed that the CoM of each link was located at the middle point of the link. CoM time-series were smoothed using a zero-lag fourth-order Butterworth filter with a cutoff frequency of 10 Hz. CoM velocity time-series were calculated using the central difference method for the filtered CoM data. For full details related to CoM estimation, see section S3 of Supplementary materials.

### Event-locked average during perturbed stance

For the perturbed condition, postural responses were characterized with respect to the perturbation-onset by event-locked average profiles of the ankle and hip joint angles as well as the position and velocity of CoM and CoP for each participant. For the event-locked average, time-series data were segmented into a number of small pieces of the data, referred to as epochs, each of which corresponds to a response to a single perturbation. Specifically, epochs from the two perturbed trails were pulled together, resulting in 40 perturbations (40 epochs) for each participant. Event-locked average time-profile was obtained by calculating the mean of the 40 epochs for each participant, where the onset of each perturbation was used as a triggering event. In addition to the participant-wise averaging, averaged profiles across participants were also computed. The event-locked average joint angles, CoM and CoP profiles were compared with the corresponding time-profiles of event-locked average responses of EMGs, ERPs and ERSPs. For our analysis, each of joint angle, joint torque, CoM, CoP, and EMG time-profiles was plotted after subtracting its mean to adjust the baseline representing the equilibrium point of each variable. Moreover, the maximum and/or the second maximum peaks of the event-locked average responses in positive and/or negative directions were detected, for which latencies from the triggering event were obtained.

### CoP analysis during perturbed stance

In the perturbed condition, the foot position shifted backward in response to every perturbation due to the backward translation of the support surface (i.e., the treadmill belt). Because the force plate and its local coordinate system were fixed in the global coordinate system independent of the moving belt, we obtained the time-series of CoP positions relative to the foot, which represents the actual postural response to the perturbation, by subtracting ankle position (that moved together with the belt) from measured CoP time-series. For full details of the CoP processing, see section S4 of Supplementary materials. The CoP time-series was then low-pass filtered in post processing using a zero-lag fourth-order Butterworth filter with a cutoff frequency of 10 Hz, while the CoP velocity time-series was obtained using the central difference method for the filtered CoP data.

To obtain a measure of postural stability throughout the duration of the perturbation, the event-locked average CoP response averaged across participants was plotted as a function of time (Fig. 1C). Moreover, they were plotted on the CoP/CoM-position vs. CoP/CoM-velocity plane (Fig. 2) as a phase plane (Bottaro et al. 2005; Bottaro et al. 2008; Asai et al. 2009).

**Figure 1.**
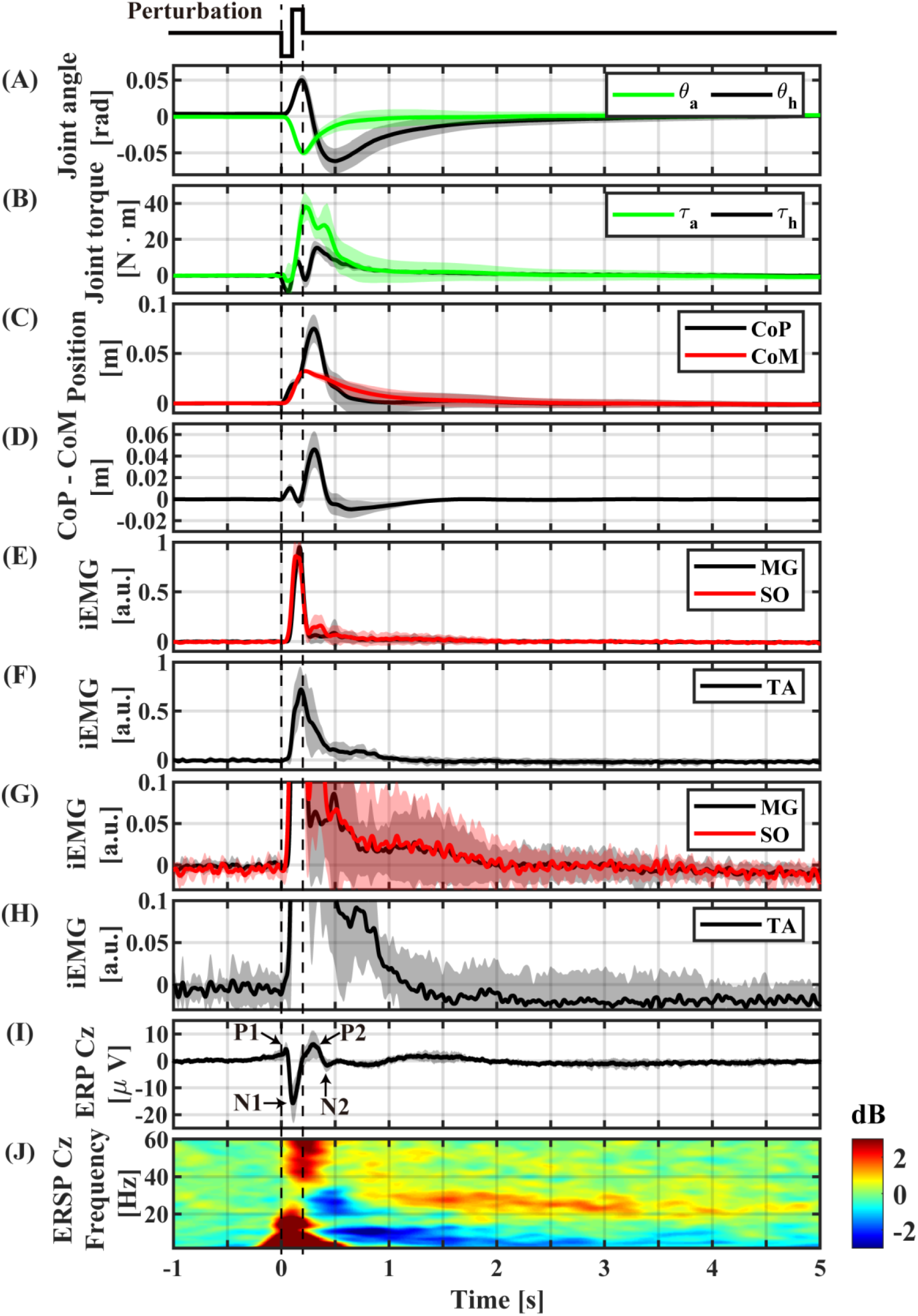
Event-locked average profiles (averaged across participants) triggered by the perturbation-onset. (A) ankle and hip joint angles, (B) joint torques obtained by the inverse dynamics analysis, (C) CoP and CoM positions, (D) CoP−CoM (difference between CoP and CoM), (E) normalized iEMGs of Medial-Gastrocnemius and Soleus, (F) normalized iEMG of Tibialis Anterior, (G) magnification of (E), (H) magnification of (F), (I) ERP of Cz electrode, (J) ERSP of Cz electrode. The light color shaded area in each of (A)-(I) is the standard deviation, representing the distribution across participants.

**Figure 2.**
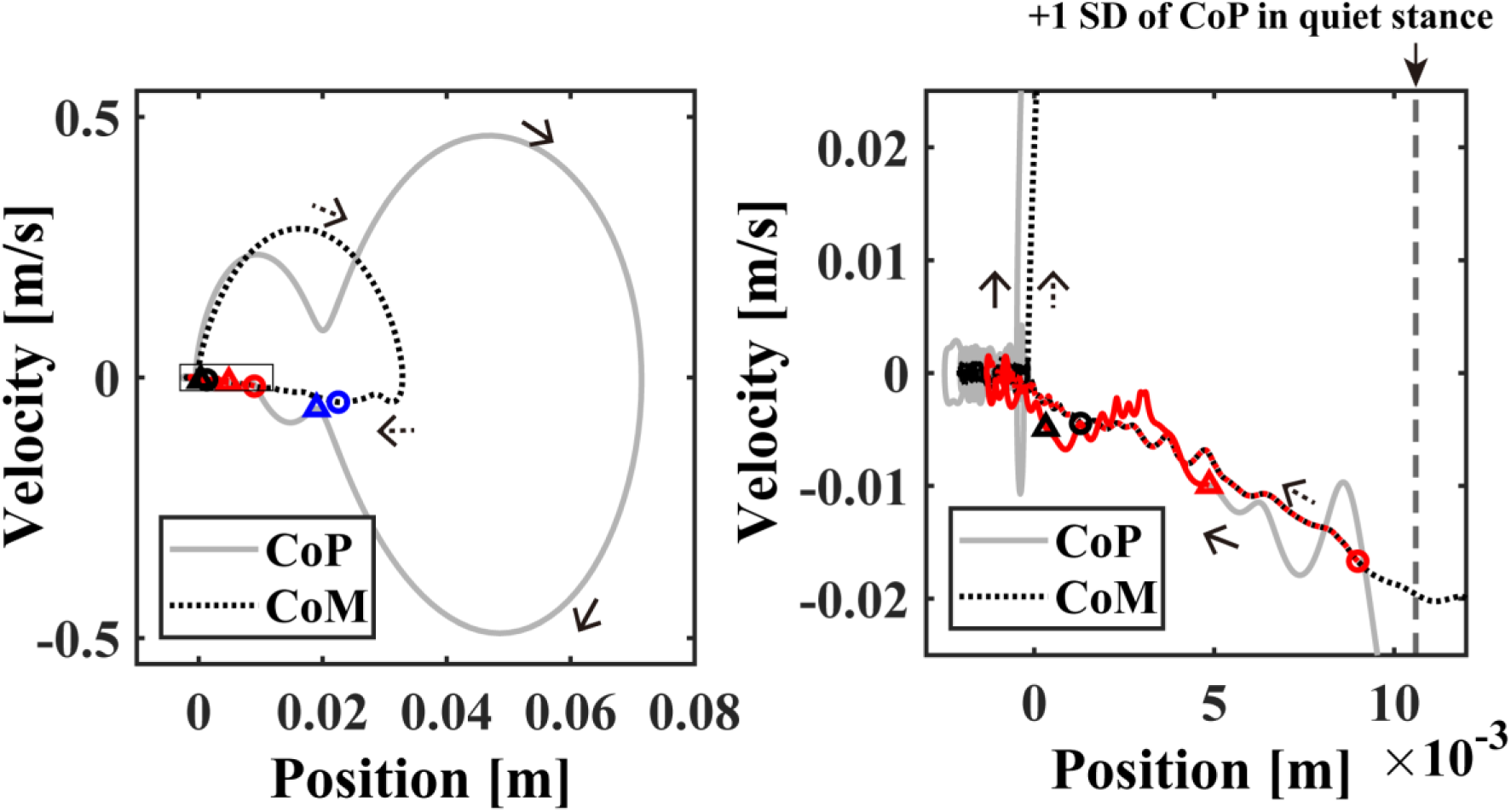
The event-locked average of CoP/CoM profile (averaged across subjects) in the CoP/CoM-position v.s. the CoP/CoM-velocity phase plane for −3 < *t* < 8 s. The small blue, red and black circles plotted on the CoM (solid-gray) trajectories in the left and/or right panels indicate the time instants of 0.5, 1.0 and 2.0 s, respectively. The triangles with the same colors indicate the corresponding time instants on the CoP (dotted) trajectories. The right panel is an enlargement of the rectangular region near the origin in the left panel. The time-duration with beta rebound (ERS) about 1.0 < *t* < 4.0 s with significantly large power was indicated by the red trajectories. The vertical dashed line in the right panel indicates the standard deviation of the postural sway during the quiet standing.

### Joint torque analysis during perturbed stance

Joint torques exerted on the ankle *τ*_a_ and the hip *τ*_h_ as the sum of passive and active torques during postural recovery responses were estimated by solving inverse dynamics based on the triple-inverted pendulum model with the foot-link fixed on the moving support-surface, the motion captured body kinematics (*θ*_a_ and *θ*_h_), the CoP positions, and the ground reaction force vectors. For full details related to the inverse dynamics analysis, see section S5 of Supplementary materials.

### EMG analysis

The EMG data recorded from MG, SO and TA muscles were processed using a zero-lag 20-450 Hz band-pass fourth-order Butterworth filter, full-wave rectified, and then low-pass filtered using a zero-lag second-order Butterworth filter with a cutoff frequency of 15 Hz (Merletti and Di Torino 1999; Yoshida et al. 2017). Because we were particularly interested in the postural dynamics in the AP-direction, the processed EMGs from the left and right legs for each muscle were averaged, which was defined as integrated EMG (iEMG) of the corresponding muscle for each participant. Then, an event-locked average of iEMG for each muscle was normalized by its maximum value using the peak of the iEMG profile for each participant. The normalized event-locked average of iEMGs were used to analyze peak latency for each muscle.

### EEG analysis

Preprocessing, denoising, and analysis of EEG signals were conducted using EEGLAB (Delorme and Makeig 2004; Loo et al. 2019). For full details about EEG preprocessing, see section S6 of Supplementary materials. Here, we present a brief summary. First, EEG data were down-sampled to 1,000 Hz. A zero-lag high-pass first-order Butterworth filter with a cutoff frequency of 1 Hz was applied (Winkler et al. 2015). We then removed data from noisy electrodes whose correlation coefficients between the surrounding electrodes were smaller than 0.8 (Bigdely-Shamlo et al. 2015). The average number of electrodes rejected in single trials was 0.28 out of 32 electrodes. We performed the artifact subspace reconstruction (ASR), which is a method for denoising EEG signals (Mullen et al. 2015). After ASR, the removed data from the noisy electrodes according to the criteria described above were replaced by the data from the surrounding electrodes using linearly spatial interpolations (Bigdely-Shamlo et al. 2015). Then, re-referencing was performed based on the average potential of all the electrodes. Independent component analysis was performed to remove the independent components (ICs) originated from EMG and electrooculograms (EOGs) activities using a method described by Bruijn et al. (2015). For single trials, the mean number of removed EMG components was 2.3, and the mean number of removed EOG components was 1.6, out of 32 ICs. After removing those artifact ICs, the remaining ICs were re-mapped onto the electrodes. The re-mapped EEG data in the time interval from 5 s prior to the perturbation until 15 s after the perturbation was then used as an epoch to analyze cortical activity.

ERP was calculated for each electrode by computing the event-locked average over all epochs (Davis et al. 1939; Quant et al. 2005). The largest and the second largest peak amplitudes in positive and negative directions for the ERP time-profile at Cz electrode were detected for each participant. Those peaks were expected to correspond to P1, N1, P2, and N2 potentials (Quant et al. 2005; Payne et al. 2018; Payne and Ting 2020a). Moreover, ERPs were plotted on the scalp as a function of time (snapshots) to characterize spatio-temporal distribution of the ERP responses (Fig. 3).

**Figure 3.**
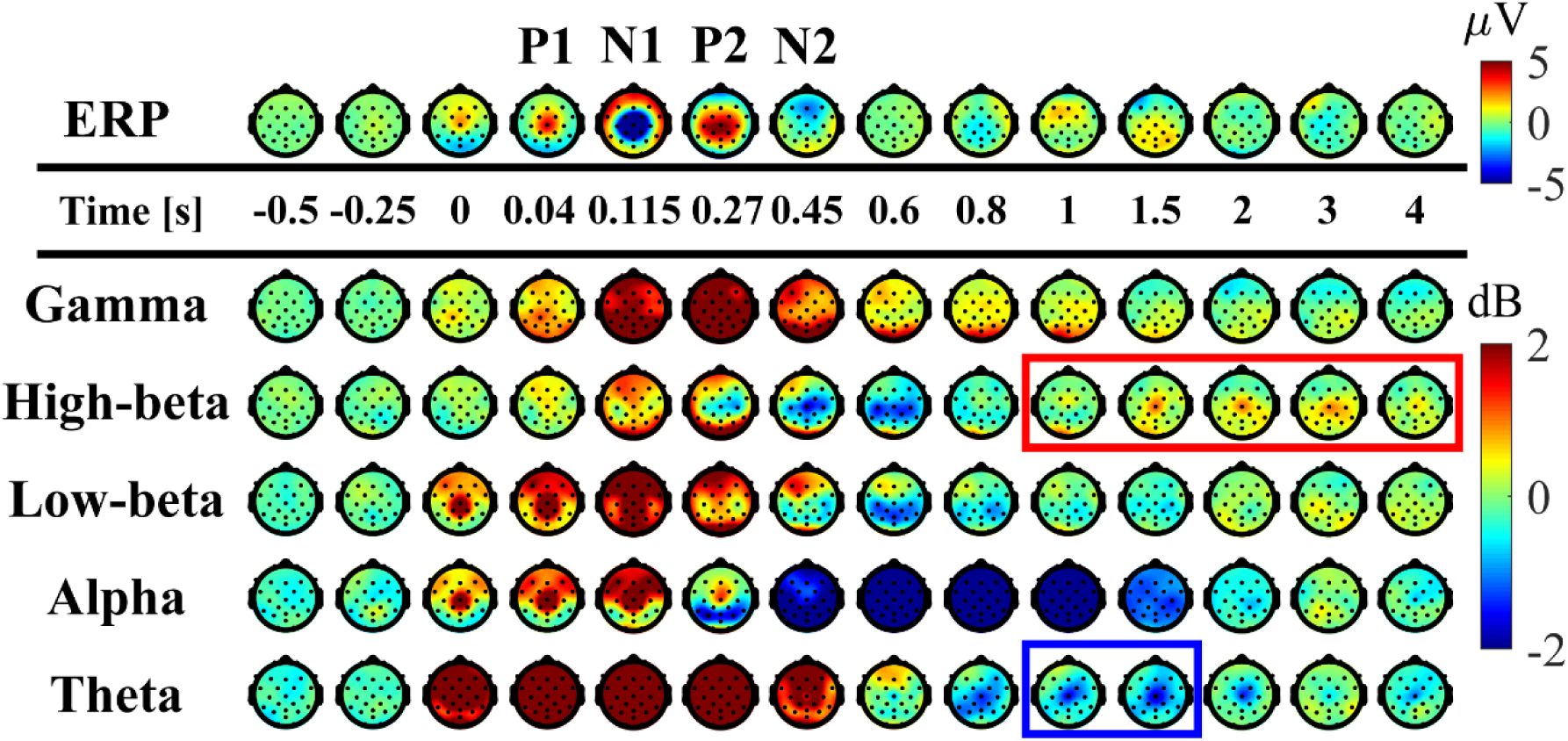
Spatial distributions of ERP and ERSP on the scalp (averaged across participants). Upper panel: time-changes in the spatial distribution of the potential on the scalp. Lower panels: time-changes in the spatial distribution of EEG-power, i.e., ERSP, for the frequency-bands of theta (3-7 Hz), alpha (8-13 Hz), low-beta (13-20 Hz), high-beta (20-30 Hz), gamma (40-60 Hz) plotted on the scalp. The scalps surrounded by the red and blue rectangulars indicate that the amplitude was significantly larger (ERS) and smaller (ERD) than the mean power prior to the perturbation.

Outcomes of time-frequency analysis using the wavelet transform for each epoch for each electrode were summarized by averaging over epochs for all perturbations, for each participant and across participants to obtain ERSPs (Makeig 1993). For full details about the wavelet transform, see section S7 of Supplementary materials. Particularly, ERSP for Cz electrode was presented by plotting wavelet coefficients as a function of time (*t*) and frequency (*f*) on the time-frequency plane. Moreover, the temporal change (snapshots) of the spatial distribution of power on the scalp using the average amplitude for each of the frequency bands at theta, alpha, low-beta, high-beta and gamma were plotted (Fig. 3).

For the ERSP on the time-frequency plane, clustered regions with enhanced (ERS) and attenuated (ERD) spectral powers were identified at each frequency band (from theta to gamma bands). To characterize such ERS and ERD at each frequency band, band-power time profiles were quantified. To this end, time axis in the time-frequency plane between −2.5 to 8 s with the perturbation-onset being at the origin was considered, and the powers at each time instant were averaged over each frequency band. See (Solis-Escalante et al. 2019) for a similar analysis. For each identified ERS/ERD, latency of the peak of the cluster were then detected using those band-power time profiles. Because prominent ERS and ERD were identified at high-beta and theta bands, which were expected to sustain for relatively long periods, time-duration of the high-beta ERS (beta rebound) and theta ERD were quantified. Time-duration of each of high-beta ERS and theta ERD was characterized for each participant using a coarse-grained-series of the band-power time profiles for the high-beta and theta bands. That is, time axis of the band-power time profiles was coarsely divided into 21 bins with time-width of 0.5 s, and the mean power over each time-bin-width was calculated to obtain the coarse-grained-series of the band-power time profiles as the function of 21 discretized latencies.

### Statistical analysis

Statistical comparisons of the time-duration of the high-beta ERS (beta rebound) and the theta ERD in ERSP were examined using multiple comparison tests for their coarse-grained-series. To this end, a distribution of mean values across participants was examined for each of 21 discretized latency, where normality of the distribution of mean values was confirmed using the Lilliefors test (Lilliefors 1967). The bin for the latency between −2.5 and −2 s was selected to represent the resting power level, which was well prior to the perturbation. The mean power of this bin was compared with that of each of the remaining 20 bins at different latencies using the Dunnett’s test (Dunnett 1955, 1964) to examine whether the power at each latency was significantly different compared to the resting power. The level of statistical significance was set at 0.05 for all tests. All statistical comparisons were performed using MATLAB Statistics Toolbox (Mathworks, USA).

## Results

### CoP during quiet stance

Postural sway during quiet stance (control condition) can be described as follows to define the stable regions for each subject. Standard deviation of sway in the AP direction was approximately 10 mm, which was about two times larger than that in the ML direction. Specifically, the average CoP sway distance in AP direction was 10.6 ± 4.9 mm and ML direction was 6.4 ± 3.8 mm. The slope of the linear regression line for the log-log plotted power spectrum of CoP in the AP direction at the low frequency regime was −1.68 ± 0.35, and that at the high frequency regime was −2.85 ± 0.48.

### Kinematic and kinetic responses to the perturbations

Kinematic and kinetic responses are summarized in Fig. 1. During support-surface acceleration phase (0 < *t* < 100 ms), the leg-link tilted forward (*θ*_a_ in Fig. 1A), i.e., the proximal end of the leg-link moved forward, due to the sudden backward shift of the feet as the distal end of the leg-link. The forward shift of the proximal end of the leg-link induced a forward shift in the distal end of the HAT-link, resulted in the backward tilt of the HAT-link relative to the leg-link (*θ*_h_ in Fig. 1A). The anti-phase movement of the ankle and hip joint angles at the initial phase of the postural response resulted in the forward shift of CoM below 0.01 m (Fig. 1C at *t* ∼ 100 ms).

During the support-surface deceleration (100 < *t* < 200 ms), inertial forces pulled the body backward in the opposite direction to that during the acceleration period. However, they were not large enough to reverse the movement directions of *θ*_a_ and *θ*_h_. Thus, the leg-link continued tilting forward, and the HAT-link continued rotating backward relative to the leg-link (Fig. 1A for 100 < *t* < 200 ms).

In the middle of the acceleration period (*t* ∼ 60 ms), SO and MG activities were initiated, and peaked at *t* ∼ 150 ms in the middle of deceleration (Fig. 1E). Those muscle activities generated plantar-flexion torque at the ankle to brake the forward tilt (Fig. 1B with the upward plantar-flexion peak of the ankle joint torque at *t* ∼ 210 ms), leading to the reverse in the rotation direction of *θ*_a_ from the negative/dorsiflexion direction toward the plantarflexion direction at the downward dorsiflexion peak at *t* ∼ 200 ms in Fig. 1A. At around the same time, TA activity reached at the peak (Fig. 1F).

The direction reversal in the leg-link (*θ*_a_) toward the upright position at *t* ∼ 200 ms initiated the backward shift of the distal end of the HAT-link, which induced the reversal in *θ*_h_ to the negative/flexion direction (the upward peak of the hip joint extension at *t* ∼ 200 ms in Fig. 1A). A contribution of the hip joint torque for this direction reversal was small, as confirmed by the small hip joint torque at *t* ∼ 200 ms in Fig. 1B.

The downward dorsiflexion peak of *θ*_a_ coincided with the peak of the CoM forward-shift at *t ∼* 200 ms in Fig. 1C. The CoP continued to move forward after the CoM peak, and peaked at *t* ∼ 300 ms. The difference between CoP and CoM (i.e., CoP−CoM) showed a qualitative agreement with the ankle joint torque (compare Figs. 1B and D).

After the dorsiflexion peak, *θ*_a_ decayed monotonously, and almost recovered the equilibrium at *t* ∼ 1.0 s. The corresponding ankle joint torque decreased to almost zero at *t* ∼ 2.0 s. The hip joint angle *θ*_h_ decreased in the negative/flexion direction until it reached the downward hip-flexion peak at *t* ∼ 500 ms, reversed again to the positive/extension direction (Fig. 1A). In contrast to the hip-extension peak, the direction reversal at the hip-flexion peak was induced by the hip joint torque peaked at *t* ∼ 330 ms in Fig. 1B. After the hip-flexion peak, *θ*_h_ and the hip joint torque decayed monotonously to the equilibrium within the interval of 2.0 < *t* < 3.0 s, which should be compared with the EEG activity lasting longer than this recovery period as shown later.

### Phase plane trajectories

Event-locked average of CoP/CoM profiles (averaged across participants) in the CoP/CoM-position vs. the CoP/CoM-velocity phase plane are shown in Fig. 2. The left panel spans the range of whole response that exhibited a perturbation-induced large excursion, whereas the right panel spans a range of postural sway during quiet stance to display the CoP/CoM responses at the very latest phase of the postural recovery. Both of CoM and CoP trajectories after *t* = 1.0 s (the red circles and triangles) approached the equilibrium posture at the origin in a linear manner (i.e., each of CoP and CoM trajectories are shaped like a line as they approach the origin) with no oscillations around the origin. The trajectory after *t* = 2.0 s (the black circles and triangles) was very close to the equilibrium, but still along the above-mentioned line, meaning that the postural state did not move randomly even within this small region near the origin. In other words, the postural recovery dynamics in this last phase still possessed strong deterministic structure that were not vanished by the event-locked averaging.

### EEG responses revealed by ERP and ERSP

Fig. 1I represents the ERP time-profile, while Fig. 1J represents the ERSP for the Cz electrode located on the primary motor area averaged across participants. In Fig. 3, the spatial distributions of ERP (top trace) and ERSP for five frequency-bands are plotted on the scalp as snapshots at several latencies for −0.5 < *t* < 4.0 s.

As in the previous study (Quant et al. 2005; Payne et al. 2018; Payne and Ting 2020a), the ERPs of P1, N1, P2 and N2 were confirmed around the Cz electrode (Fig. 1I and 3-top). Those ERP potentials could be decomposed into ERS at the gamma, low-beta, alpha and theta frequency-bands that appeared all together within the initial phase before *t* ∼ 300 ms after the perturbation-onset (Fig. 1J and Fig. 3).

Latency and peak amplitude of the P1 potential was 41 ± 18 ms and 5.1 ± 2.3 μV, respectively, which was immediate before the onset of SO, MG and TA activations. The P1 was mainly composed of ERSs at low-beta and alpha bands, each of which was spatially localized around the Cz electrode (Fig. 3 at 40 ms).

Latency and peak amplitude of the N1 potential was 116 ± 22 ms and −17.2 ± 6.2 μV, respectively, which was immediately after the onset of support-surface deceleration, and followed a few tens of milliseconds by the peaks of SO, MG and TA activities (Fig. 1E-F). The N1 at the Cz electrode location was composed of the low-beta ERS that peaked at 94 ± 20 ms, alpha ERS that peaked at 92 ± 26 ms, and theta ERS that peaked at 124 ± 30 ms with a minor contribution of the gamma ERS (Fig. 1J). The low-beta ERS and alpha ERS for N1 were part of (continuation of) the corresponding ERSs that contributes to P1. Note that the ERS contributing to P1 and N1 potentials was absent specifically for the high-beta band. The low-beta ERS and theta ERS as well as the gamma ERS for N1 were spatially distributed over the scalp, but the alpha ERS was localized around Cz electrode (Fig. 3).

Latency and peak amplitude of the P2 potential was 264 ± 49 ms and 8.5 ± 3.6 μV, respectively, which was a few tens of milliseconds after the termination of the support-surface deceleration, immediately after the end of large activations of SO, MG, middle of the large TA activation. It also corresponded to the peak of forward-shift of CoP (Fig. 1C). Moreover, it coincided in time with the largest peak of the hip extension torque (Fig. 1B). P2 was mainly composed of the gamma ERS peaked at 252 ± 106 ms for Cz (Fig. 1J). The gamma ERS for P2 was spatially distributed over the scalp. Although P1 and P2 were contributed from the theta ERS distributed over the scalp, the appearance of the theta ERS for P1 and P2 was due to the low resolution (the uncertainty principle) of the wavelet at the low-frequency bands.

During the postural recovery phase (300 < *t* < 600 ms), ERD at high-beta band (high-beta ERD) was observed dominantly at Cz (Fig. 1J) and weakly at C_3_ and C_4_ (Fig. 3). The peak latency of the high-beta ERD was 450 ± 83 ms for Cz, which was roughly coincided with the N2 potential (Fig. 1J). Moreover, the high-beta ERD coincided also with the time instant when the CoP on the way back to the upright position caught and surpassed the CoM that preceded the CoP until this point (Fig. 1C), which altered the sign of CoP−CoM, representing the ankle torque, from positive to negative (Fig. 1D). Note that the positive and negative ankle torques represent, respectively, pulling the forward-tilted posture backward and braking the backward moving posture. This ankle torque was mostly generated by body mechanics induced by the relative positioning between CoP and CoM, which could be confirmed by the fact that the large EMG responses of SO, MG and TA were already in the late of their decreasing phases. However, those muscle activities also contributed to the stabilization (Figs. 1B, 1G and 1H). In this way, the high-beta ERD did not necessarily appear in the middle of the large EMG responses, but it appeared at the tail of the large responses of SO and MG. Since the large activation of TA was delayed relative to those of SO and MG (about 50 ms difference in the peak latencies), the high-beta ERD overlapped with the late phase of the large TA activation that brakes the backward moving posture.

For latencies longer than 1.0 s, we showed two event-related clusters of powers in ERSP (Fig. 1J). One was the ERS at high-beta band (high-beta ERS), which appeared after the high-beta ERD. The high-beta ERS lasted over a relatively long duration with its peak latency located at 3.27 ± 1.35 s. The high-beta ERS was observed mainly at Cz electrode (Fig. 3). Fig. 4 shows time evolution of the high-beta ERS at Cz electrode. Distributions of the powers of high-beta band at discretized bins were normal, except the bins at [-0.5, 0] s and [-5.5, 6.5] s. The averaged powers across participants at the high-beta band in the latency of 1 < *t* < 4 s, for about 3 s, were significantly greater than the resting potential (*p* < 0.05).

**Figure 4.**
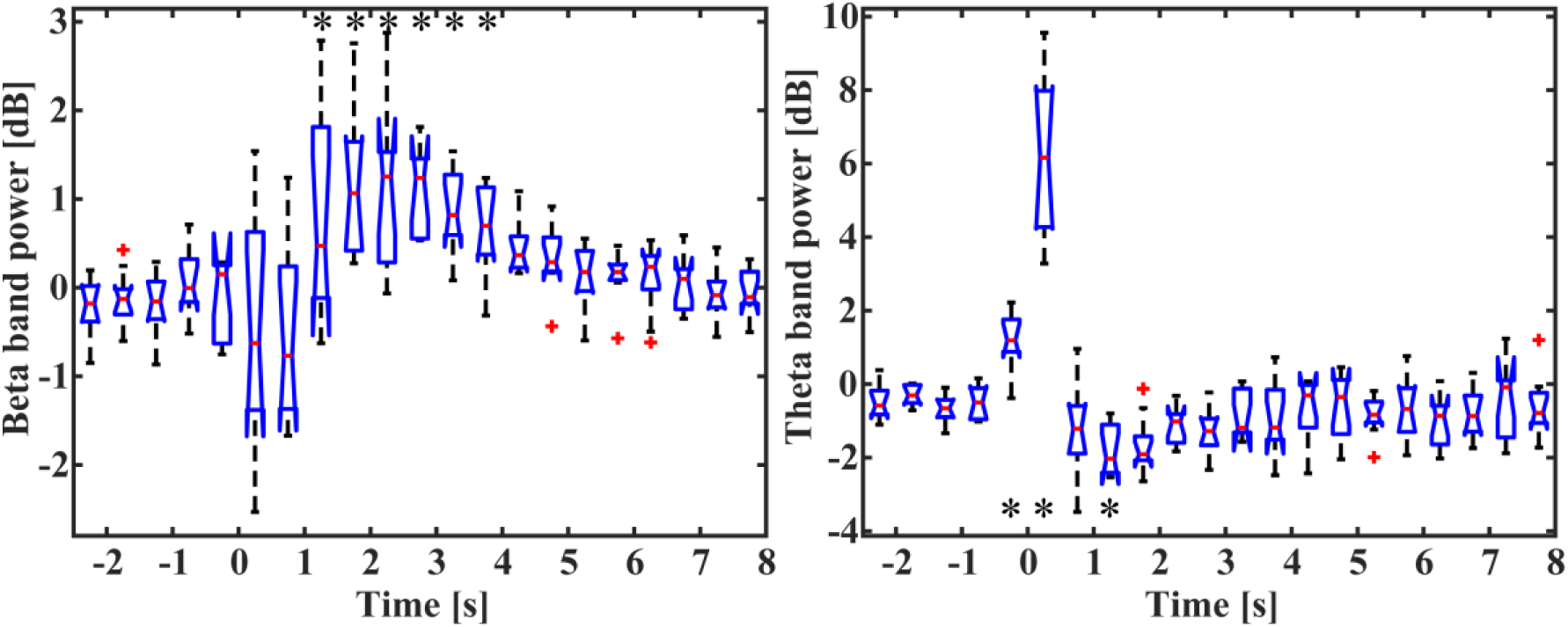
Time-changes in the average powers of high-beta-band (left) and theta-band (right). *t* = 0 corresponds to the perturbation-onset. Average power values were calculated for each participant for −2.5 < *t* < 8 s for every 0.5 s interval representing each box of the plots. The powers of the boxes with the symbol * were significantly larger and smaller than the resting potential (*p* < 0.05, Dunnett’s test), respectively, for the high-beta and theta bands.

During the appearance of the high-beta ERS, the CoP and the CoM became close to each other in the very late phase of the recovery process (Figs. 1A and C). This could also be confirmed by Fig. 2 that shows the trajectories of postural response in the CoP/CoM-CoP/CoM velocity phase plane. Particularly, the trajectories within the period of time when the high-beta ERS was significantly large is indicated by the red curve segments in the phase plane of Fig. 2, by which we could observe that the postural state during the appearance of the high-beta ERS was located in the area very close to the range of postural sway during quiet stance. However, as described above, the postural state during the high-beta ERS did not move randomly, but still it moved mostly deterministically along the above-mentioned line. Comparison between ERSP in Fig. 1J and enlarged EMGs in Figs. 1G and H showed that activations of SO and MG muscles were still larger than those during quiet stance for the early period (1 < *t* < 3 s) of the high-beta ERS. However, activations of all muscles were almost the same as those during quiet stance for the later phase (3 < *t* < 4 s) of the high-beta ERS. That is, the high-beta ERS in its later phase was generated with almost no perturbation-induced muscular activations.

In addition to the high-beta band, ERD at alpha and theta bands were also observed. As shown in Fig. 3, the alpha ERD appeared at the occipital region for the early phase (*t* ∼ 270 ms), and then expanded to the whole head, with the latency 0.83 ± 0.17 s at Cz. The time interval of the alpha ERD coincided with the period of time when the sign of CoP−CoM was negative for braking the backward-moving posture (Fig. 1D). Then, the theta ERD appeared specifically around Cz with its latency of 1.88 ± 0.74 s. Fig. 4-right quantifies the time-range of the theta ERD. Distributions of the powers of theta band at discretized bins were normal, except the bins at [1.5, 2] s and [3.5, 4] s. The averaged powers across participants at the theta-beta band at Cz electrode in the latency bin of [1, 1.5] s was significantly smaller than the resting potential (*p* < 0.05). However, although no statistical significance was obtained, the theta ERD also exhibited a long-lasting property as in the high-beta ERS, which could be visually confirmed in Fig. 1J with the blue-colored ERD at theta band until the latency of 3.0 s.

## Discussion

In this study, we examined the postural recovery process during upright stance in response to the support-surface impulsive perturbation. Slower time-scale of mechanical dynamics during postural recovery of the perturbed posture, compared to movements of the upper extremities, prompted our analysis of the temporal profile of cortical activations during postural stabilizing process. It was shown that neural responses for re-stabilizing upright posture lasted over a relatively long periods of time (i.e., few seconds), which is consistent to our hypothesis. Specifically, we identified a novel type of high-beta band cortical synchronization activity (high-beta ERD and high-beta ERS or beta rebound) localized around Cz electrode over the primary motor area. The appearance of beta ERD and ERS is consistent to the response profile of the upper extremities, while the responses were much shorter and typically lasting only for a few hundred milliseconds or 1 second in the longest case (Engel and Fries 2010; Pfurtscheller et al. 1996; Pfurtscheller and Lopes da Silva 1999). However, the temporal configuration (temporal location) of the beta ERS relative to the kinematic and kinetic responses of the perturbed stance was different qualitatively from those for the upper extremities. That is, the high-beta ERS for the perturbed stance started when the postural recovery process was still present, and it was long-lasting for about 3 s along with a residual motion during the postural recovery (Fig. 1J and Fig. 3). This is in stark contrast to the beta ERS for the upper extremities that started after movements are terminated.

### Responses in the initial phase of the perturbation

Before discussing our main finding, which occurred in the later part of the postural response, reflexive cortical responses during the initial phase of the postural recovery process within about 300 ms after the perturbation-onset are summarized and discussed first. We confirmed P1, N1, and P2 peaks of the ERPs (Fig. 1J and Fig. 3), consistent with the previous studies (Quant et al. 2005; Payne et al. 2018; Payne and Ting 2020a). Specifically, the P1 potential (with the peak latency of 40 ms) was composed of the low-beta and alpha ERSs that were located at Cz electrode, and occurred immediately before the onset of SO, MG and TA activations (Fig. 1E-F). The short latency of P1 (slightly less than 50 ms) is similar to that of vestibular-evoked potential (Todd et al. 2014), which might trigger centrally the initial muscular responses.

Additional recruitment of the ERS components at the theta band altered the P1 potential into the N1 potential with the peak latency of 120 ms as reported in the previous studies (Mierau et al. 2017; Varghese et al. 2014). It has been considered that N1 represents neural processing of sensory information necessary for coordinating reactive balance responses (Dietz et al. 1984, 1985; Dimitrov et al. 1996; Quant et al. 2004; Staines et al. 2001; Payne et al. 2018; Payne and Ting 2020a). Moreover, it might represent an error signal for postural instability (Adkin et al. 2006, 2008; Mochizuki et al. 2009; Payne et al. 2019). Such sensory information processing might contribute to enhancing the muscle activations of SO and MG that were initiated by the spinal stretch reflexes, since the N1 potential was followed by large activations of SO and MG (Fig. 1E). Moreover, large response of TA muscle peaked at 200 ms, which overrode a possible action of reciprocal inhibition from the SO/MG activations. This might be initiated centrally by the N1 potential to stiffen the ankle joint via counteracting the SO/MG-induced plantar-flexion ankle torque that peaked at 150 ms. A recent study reported that N1 amplitudes alter depending on postural balance functionality, i.e., larger N1 reported for individuals with lower balance ability and smaller N1 responses for those with better balance (Payne and Ting 2020a). Taken together, these results imply that N1 expresses a supraspinal modulation of spinal reflexes.

The P2 potential was predominantly composed of the gamma ERS, which might represent a neural detection of postural instability (Slobounov et al. 2005). It was followed by the largest peak of hip extension torque, implying that the P2 potential might be associated with the generation of extension torque at the hip joint (Figs. 1B and J). Further examinations with EMGs of hip joint muscles are required to validate the active contribution of the P2 to the control of hip joint movement. In this way, we confirmed typical ERPs with short latencies that have been examined so far during perturbed stance. Existence of those ERPs despite the differences in detailed experimental setup and protocol indicates that they are robust and stereotypical.

### Desynchronization of cortical activities at the end of the initial reflexive phase

In this study, the high-beta ERD was observed predominantly around the primary motor area. It peaked at 450 ms, roughly coincided with the N2 potential. In general, high-beta ERDs have been identified for motor tasks other than the upright stance, in which they appear at the beginning of, or sometimes slightly prior to, the movement and sustain for the duration of the movement (Jurkiewicz et al. 2006; Neuper and Pfurtscheller 1996; Pfurtscheller et al. 1996; Pfurtscheller and Lopes da Silva 1999; Rossiter et al. 2014). Specifically, it was shown to appears continuously during sustained voluntary movements of hand and finger (Erbil and Ungan 2007; Nakayashiki et al. 2014). A recent study showed that a smooth execution of a voluntary movement requires a decrease in the power of beta band (Heinrichs-Graham and Wilson 2016), suggesting that beta ERDs are associated with the preparation of actions and related information processing to facilitate a motor execution. Note that beta ERDs appear even in passive (non-voluntary) motor tasks (Vinding et al. 2019) and also during the Go/NoGo tasks regardless of the motor selection, i.e., either Go or NoGo responses (Alegre et al. 2004; Wu et al. 2019). That is, beta ERD (and ERS) is not necessarily accompanied with actual motor executions, but it represents information processing of afferent sensory signals as well as the decision-making process (Jurkiewicz et al. 2006; Alegre et al. 2004; Wu et al. 2019). The high-beta ERD in this study appeared when the CoP recovering toward the upright position caught and surpassed the CoM, after which CoP and CoM became close to each other. It also coincided with the period of time when the muscle activations became much smaller than those in the initial phase of the postural response for *t* < 300 ms. This seemingly contradicts a typical appearance of beta ERD because the high-beta ERD in this study was not accompanied with the major muscle activations for *t* < 300 ms. However, the major muscle activations with the large postural response for *t* < 300 ms could be considered as a stereotypical sequence of reflexes both at spinal and cortical levels, and the postural control after the initial phase might require sophisticated information processing to achieve postural recovery, which might be represented in the high-beta ERD in this study. For example, the high-beta ERD after the initial postural response could represent a state of preparing for possible actions after the perturbation, such as a stepping response in case that the perturbation is larger than a specific threshold. Specifically, this can be achieved by inhibiting the reflexive muscular reactions during the initial phase of the postural response and by facilitating the cortico-spinal excitability (Solopova et al. 2003). Those interpretations are consistent with the recent report on N2 potential reflecting specific cortical responses to balance perturbations during upright stance, which are affected in PD patients (Payne and Ting 2020b).

It is of interest to note that slightly before the high-beta ERD (*t* ∼ 270 ms), the alpha ERD appeared at the occipital region, which might correspond to the previously reported ERD at mu band (Jurkiewicz et al. 2006), and then expanded throughout the cortex until about *t* ∼ 1 s (Fig. 3). Then, the theta ERD appeared specifically around Cz with the latency of *t* ∼ 800 ms lasting for at least 0.5 s (Fig. 4-right). During the time interval with the alpha ERD and theta ERD, the CoM was located close to or slightly ahead of the CoP, while the body was recovering toward the upright position. Although no statistical significance was obtained, the theta ERD also exhibited a long-lasting property as in the high-beta ERS until the latency of *t* ∼ 3 s. The appearance of movement-related theta ERD is previously unreported to our best knowledge. Because it sustained for a long period in parallel with the beta rebound, it might play a coordinated role with the beta rebound in the sensory information processing.

### Long-lasting high-beta rebound

A high-beta ERS, also known as beta rebound, typically appears after completion of a movement, and often following the corresponding beta ERD (Jurkiewicz et al. 2006; Neuper and Pfurtscheller 1996; Pfurtscheller et al. 1996; Pfurtscheller and Lopes da Silva 1999). Consistently in this study, the high-beta ERS also appeared after the high-beta ERD. However, it began before the postural recovery was completed (Figs. 1A and J), which is different from previous studies examining upper extremities. The simplest interpretation is that the major part of the postural response was completed within the initial phase of the recovery process, and that the small residual response that remained during the beta ERS is negligible. If this is the case, we could consider that the high-beta ERS in this study was also initiated after the movement as in the previous studies for the upper extremities, implying the same functional roles expressed by the high-beta ERS in this study and previous upper extremity studies. That is, the cortical beta activity in humans represents a raise in the cortical activity levels that suppress voluntary movement (Gilbertson et al. 2005; Pastötter et al. 2008; van Wijk et al. 2009), while actively promoting postural maintenance (Gilbertson et al. 2005). The promotion of postural maintenance may be achieved through an up-regulation of relevant sensory inputs during and immediately after bursts of beta activity (Androulidakis et al. 2006; Gilbertson et al. 2005; Lalo et al. 2008). Indeed, beta ERS during finger movement is reduced in patients with impaired afferent sensory information processing due to Parkinson’s disease (PD) (Vinding et al. 2019). Moreover, ischemia in the peripheral afferent reduces the power of beta rebound (Cassim et al. 2001). As in the beta ERD, the beta ERS appears not only after actual motor execution, but also after a motor imagery and/or after NoGo response in Go/NoGo tasks (Pfurtscheller and Lopes da Silva 1999; Wu et al. 2019). Moreover, the beta ERS in PD patients is attenuated (Wu et al. 2019). Those evidences suggest that the beta ERS represents a processing of afferent sensory information (Jurkiewicz et al. 2006) and a decision-making process (Alegre et al. 2004; Wu et al. 2019) in combination with the beta ERD.

An alternative and novel interpretation is required if the small residual postural dynamics during the beta ERS are considered not negligible. In this case, we should conclude that the beta ERS appeared before the perturbation-induced movement was completed. This is in stark contrast to the beta ERS for the upper extremities, implying that we found a novel process with the beta rebound before movement completion. This novel interpretation can be supported by the fact that the CoP/CoM state points during the beta ERS still exhibited strong deterministic structure that was not vanished through event-locked averaging (Fig. 2-right). Moreover, the small residual error that remained during the beta ERS, which is the one we neglected for interpreting the onset of the beta ERS in the discussion above, would be the only possible cause that could drive the long-sustained high-beta ERS for about 3 s. In other words, if the residual error were negligibly small, it could not drive the high-beta ERS for the long time, and thus, the high-beta ERS should have completed in a shorter duration than that sustained over the course of three seconds. Thus, it turns out that the small residual postural dynamics converging to the upright equilibrium might not be simply an “almost equilibrated state”, but they should still be taken care of for achieving postural stabilization, which requires a certain amount of information processing of afferent sensory signals, leading to the long-lasting high-beta ERS.

We propose that the long-lasting high-beta ERS at the late phase of postural recovery might be associated with the recently proposed intermittent control during human quiet standing (Bottaro et al. 2005; Bottaro et al. 2008; Asai et al. 2009). For a comprehensive summary of the model, see Nomura et al. (2020). In short, the intermittent control model assumes a postural-state-dependent switching between inactivation (off) and activation (on) of the neural feedback controller with time-delay, which plays a critical role for stabilizing the upright posture. Specifically, the model assumes a line-shaped switching boundary passing through the origin that separates a state space of an inverted pendulum (representing the upright posture) into an off-region and an on-region. The feedback controller is deactivated if the state point is in the off-region and, otherwise, activated in the on-region, where appropriate actions of the switch-off are crucial for the stability. Typical dynamics of the intermittent control model (Bottaro et al. 2005; Asai et al. 2009) are quite similar to the experimental postural response at the late phase of the perturbed stance near the quiet standing region in Fig. 2-right of this study, where both the model and experimental subjects exhibited a seemingly overdamped-system-like non-oscillatory linear trajectory toward the equilibrium posture in the very late phase of the postural recovery. This seemingly overdamped-system-like trajectory in the intermittent control model was indeed achieved by the off-on chattering that takes place typically along the line-shaped switching boundary (see Fig. 9 in Bottaro et al. 2005 or Fig. 3A in Asai et al. 2009), not by the overdamped impedance controller. Although the intermittent feedback control is involuntary for the automatic postural stabilization, it is highly probable that the decision-making processes for selecting on- or off-switching might generate high-beta ERD and ERS as in decision-making processes for selecting Go or NoGo actions in Go/NoGo tasks (Alegre et al. 2004; Swann et al. 2009; Zhang et al. 2008; Wu et al. 2019).

## Acknowledgements

We thank Dr. Hikaru Yokoyama for an initial guidance on performing EEG measurements and related signal processing methodologies.

## Grants

This study was supported by the following grants from the Ministry of Education, Culture, Sports, Science and Technology (MEXT)/Japan Society for the Promotion of Science (JSPS) KAKENHI: 20H05470 (T.N), 19H04181 (T.N), 17K13016 (Y.S), and 20K19412 (M.M), and by the Uehara memorial foundation (T.N).

## Disclosures

No conflicts of interest, financial or otherwise, are declared by the authors.

## Endnote

At the request of the authors, readers are herein alerted to the fact that additional materials related to this manuscript may be found at https://doi.org/10.5281/zenodo.3955496. These materials are not part of this manuscript, and have not undergone peer review by the American Physiological Society (APS). APS and the journal editors take no responsibility for these materials, for the website address, or for any links to or from it.

## Author Contribution

A.N. and T.N. conceived and designed research; A.N. and Y.S. performed experiments; A.N. analyzed data; Y.S and M.M advised on experimental and analytical methodologies; A.N. and T.N. interpreted results of experiments; A.N. prepared figures; A.N. drafted manuscript; Y.S., M.M. and T.N. edited and revised manuscript; A.N., Y.S., M.M. and T.N. approved final version of manuscript.

## Notes

https://doi.org/10.5281/zenodo.3955496

### Competing Interest Statement

The authors have declared no competing interest.

